## Supplementary figures and images for "Comparing the Performance of mScarlet-I, mRuby3, and mCherry as FRET Acceptors for mNeonGreen"

### Supplemental Figures

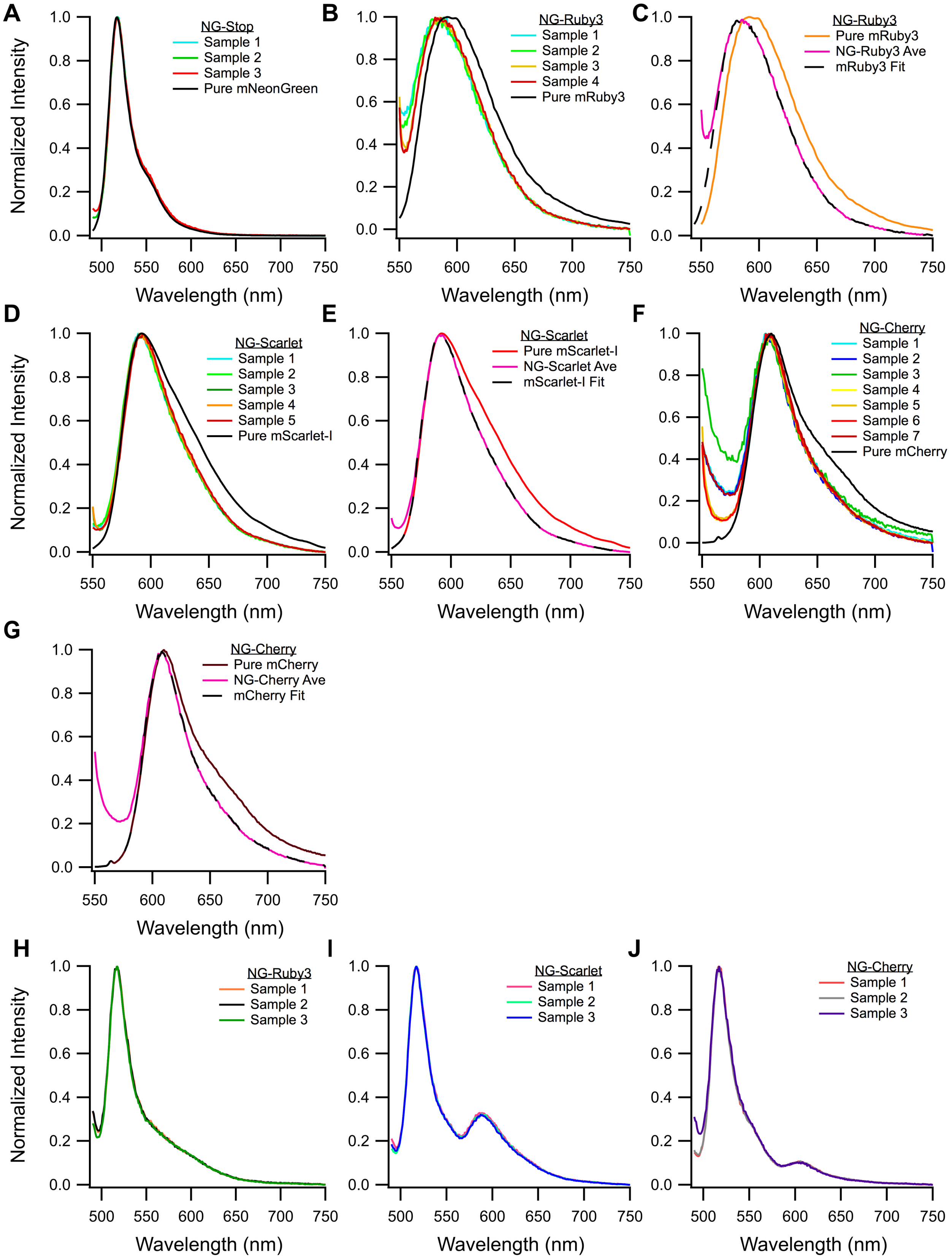

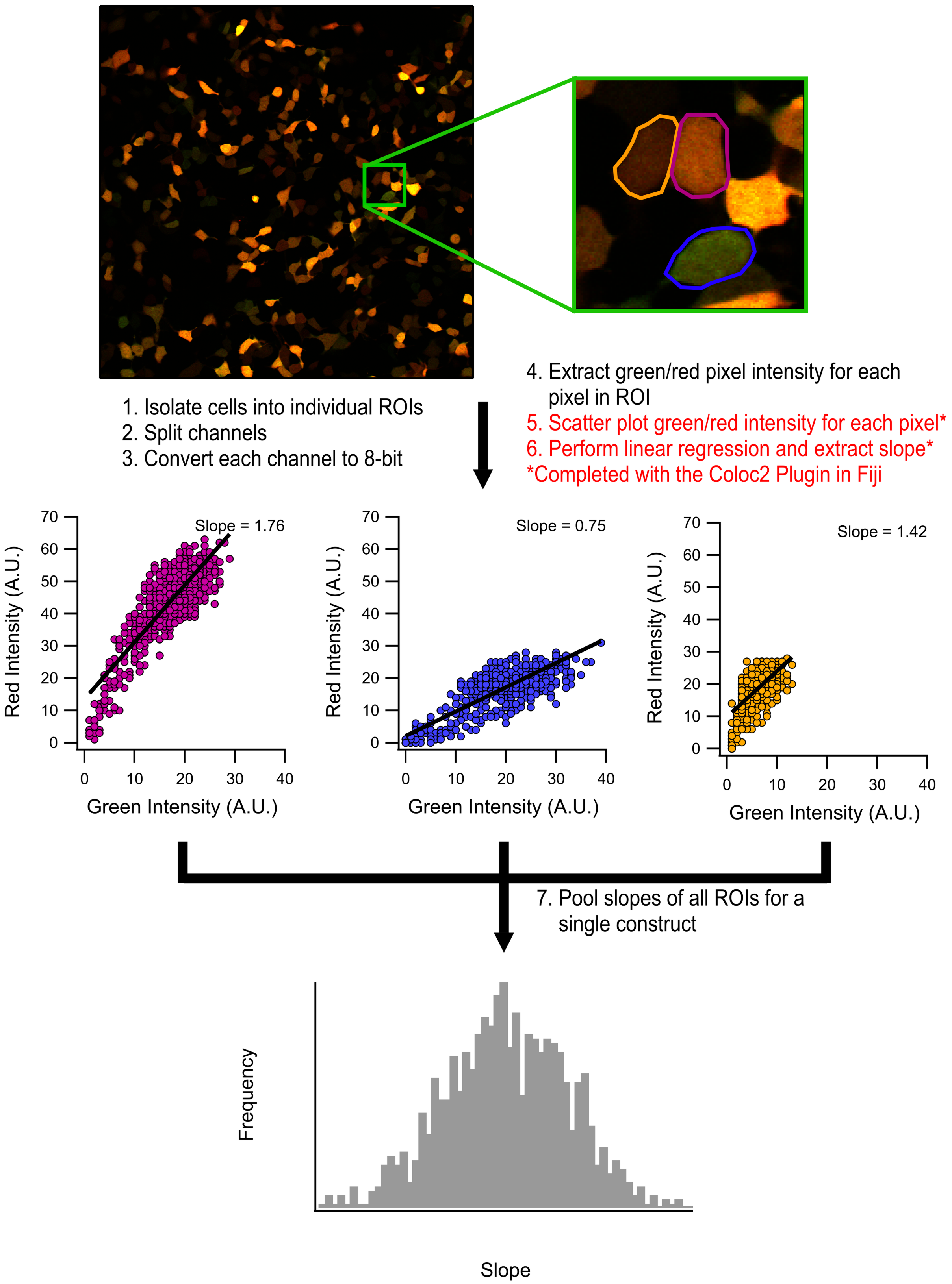

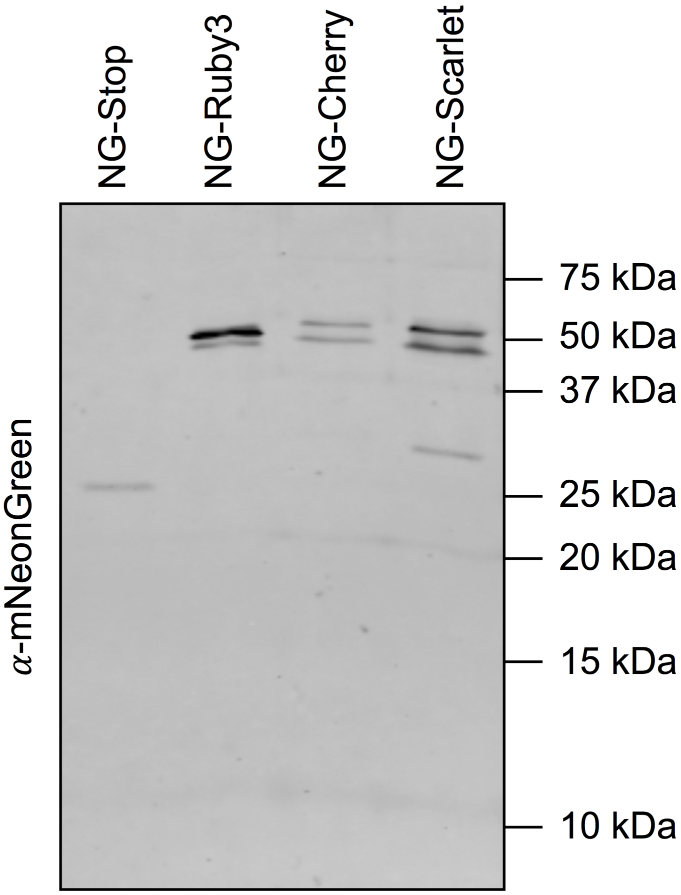

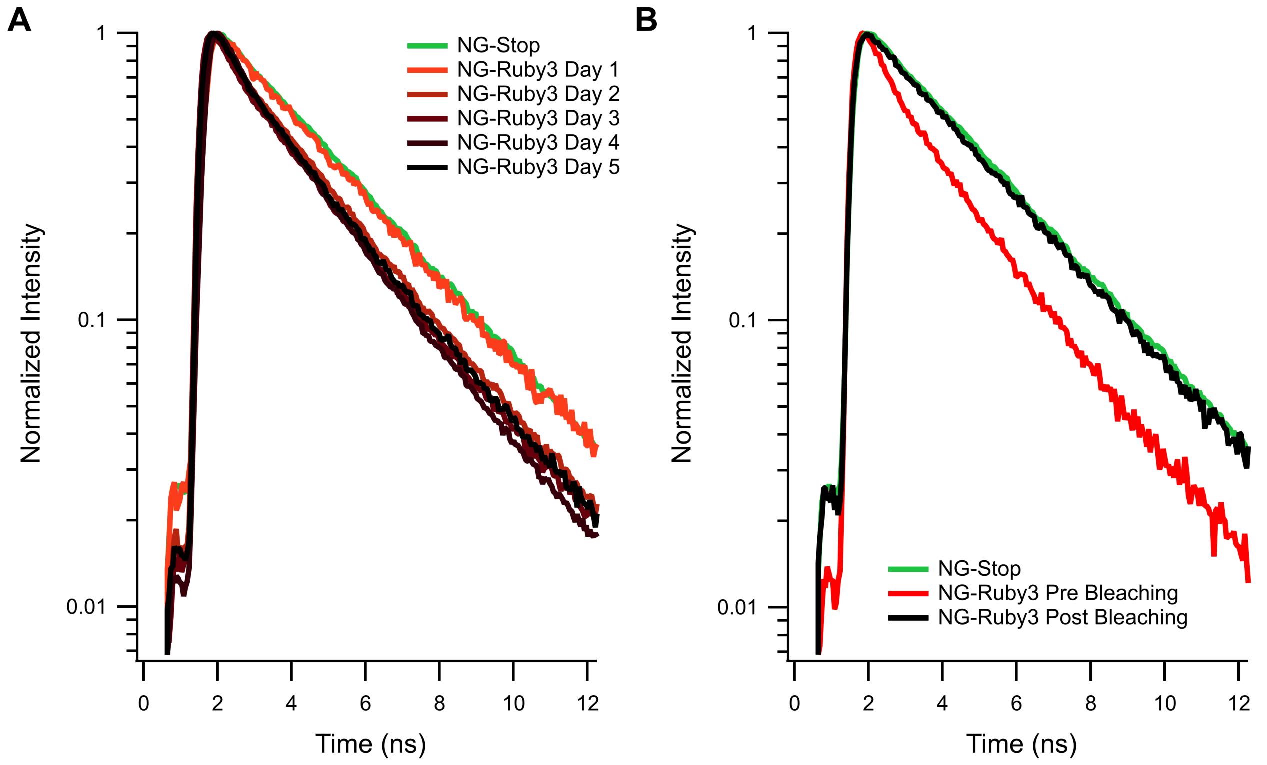
